# The presynaptic active zone protein Bassoon as a marker for synapses between Type lll cells and afferent nerve fibers in taste buds

**DOI:** 10.1101/2022.04.06.487279

**Authors:** Rio Ikuta, Shun Hamada

## Abstract

Taste buds are receptor organs for gustation. Two types of taste receptor cells have been identified in taste buds: Type II and Type III cells. Type III cells connect with afferent fibers through conventional chemical synapses. In the present study on taste buds, we used immunocytochemistry to examine the distribution pattern of Bassoon, a scaffolding protein of the cytomatrix at the active zones of conventional synapses. Bassoon was predominantly detected as small puncta in Type III cells. Bassoon-immunoreactive puncta were observed in proximity to or partially overlapping with intragemmal nerve fibers. Immunoelectron microscopy showed Bassoon at the active zones of the conventional synapses of Type III cells. The present results demonstrated that Bassoon is a marker for synapses between Type III cells and afferent fibers in taste buds.

## Introduction

Taste buds are receptor organs for gustation and are mainly present in the epithelium of the papillae of the tongue. Each taste bud consists of 50-100 cells that are classified into four types (Types l, ll, lll, and IV) based on their morphological characteristics (Murray, 1993; Murray et al., 1969). Type II and III cells are directly involved in the transduction of different taste modalities and activate afferent nerve fibers (review, Finger and Barlow, 2021).

Type II and III cells both form synapses with afferent gustatory nerve fibers (Wilson et al., 2021; Yang et al., 2020). However, the structures of synapses in Type ll and III cells are markedly different from each other. The synapses between Type III cells and afferent nerve fibers are conventional synapses that show the accumulation of synaptic vesicles with presynaptic densities (Murray et al., 1969; Royer and Kinnamon, 1991; Takeda, 1976; Takeda and Hoshino, 1975; Yang et al., 2020). In contrast, the synapses of Type II cells lack synaptic vesicles and release ATP as a neurotransmitter through CALHM1/3 ATP channels (Ma, et al., 2018; Romanov, et al., 2018). Thus, it has been proposed that synapses of Type II cells should be referred to as “channel synapses” (Taruno, et al., 2020).

Reliable markers for synapses at the light microscopic level greatly contribute to the study of synapse formation in various nervous systems. Synaptic markers are important for the study of synapses in taste buds because they are continuously formed throughout life as taste bud cells are renewed (Finger and Barlow, 2021). CALHM1 has been suggested as a presynaptic marker for channel synapses between Type II cells and nerve fibers (Kashio, et al., 2019;Romanov, et al., 2018), whereas there is currently no useful marker for conventional synapses between Type III cells and afferent nerve fibers. To date, the localization of several synaptic proteins in conventional synapses has been investigated in taste buds (Asano-Miyoshi, et al., 2009;Finger, et al., 1990;Kohno, et al., 2005;Kotani, et al., 2013;Pumplin and Getschman, 2000;Yang, et al., 2000;Yang, et al., 2007;Yang, et al., 2004). However, none of these synaptic proteins are markers for the conventional synapses of Type III cells because they did not exclusively localize to synaptic sites in taste buds.

In the present study, we focused on Bassoon, a scaffolding protein of the cytomatrix at the presynaptic active zone (tom Dieck et al., 1998), which is involved in the assembly of active zones, the priming of synaptic vesicles, and the localization of voltage-gated Ca2+ channels to the active zones (review, Gundelfinger et al., 2015). Bassoon selectively localizes to the active zones of conventional and ribbon synapses in the central and peripheral nervous systems (Gundelfinger et al., 2015), but has not yet been investigated in taste buds. Therefore, we herein examined the distribution pattern of Bassoon in taste buds by immunohistochemistry using light and electron microscopy.

## Materials and methods

Eight-week-old male ICR mice were obtained from Japan SLC Inc. (Shizuoka, Japan). Mice were housed with a standard laboratory diet and water *ad libitum*. All experimental procedures were performed with the approval of the Institutional Animal Care and Use Committee of Fukuoka Women’s University.

Mice were deeply anesthetized with sodium pentobarbital and then perfused through the ascending aorta with saline followed by fixatives for 10 minutes. As fixatives, we used 2% paraformaldehyde in 0.1 M phosphate buffer (PB, pH 7.3) for light microscopy and periodate-lysine-paraformaldehyde fixative for immunoelectron microscopy (McLean & Nakane, 1974). After perfusion fixation, the circumvallate papillae were removed and postfixed in the same fixative at 4°C for 2 hr. Tissues were then immersed in 25% sucrose in 0.1 M PB at 4°C overnight for cryoprotection and frozen in embedding medium with hexane cooled by dry ice. Frozen tissues were stored at −80°C for later analyses.

### Immunofluorescence

Tissues were cut into 7-µm-thick sections on a cryostat and collected on slides. Sections were rinsed in phosphate-buffered saline (PBS) containing 0.3% Triton X-100 (PBST), and then treated with a blocking reagent (Blocking One, Nacalai Tesque, Japan) containing 3% M.O.M Blocking Reagent (Vector, CA, USA) for 1 hr. After a brief rinse with PBS, sections were incubated with a mixture of the primary antibodies diluted with PBST containing 5% Blocking One at 4°C overnight. The primary and secondary antibodies used in the present study are listed in Table 1. After an incubation with the primary antibodies, sections were rinsed with PBST and PBS. Sections were incubated with appropriate secondary antibodies labeled with Alexa 488 at room temperature for 1 hr. After rinses with PBST and PBS, the other secondary antibodies labeled with Alexa 594 were applied to the sections for 1 hr. Regarding triple immunolabeling, sections were sequentially incubated with appropriate secondary antibodies labeled with Alexa 488, Alexa 555, and Alexa 647 for 1 hr each. All secondary antibodies (Table 1) were used at a dilution of 1:1000. Sections were rinsed with PBST and PBS, incubated with DAPI, and then coverslipped with mounting medium. Controls were prepared by omitting the primary antibodies. Images were collected under a confocal laser microscope (C2 plus, Nikon, Japan). The contrast and brightness of images were adjusted with Adobe Photoshop.

**Table 1.**
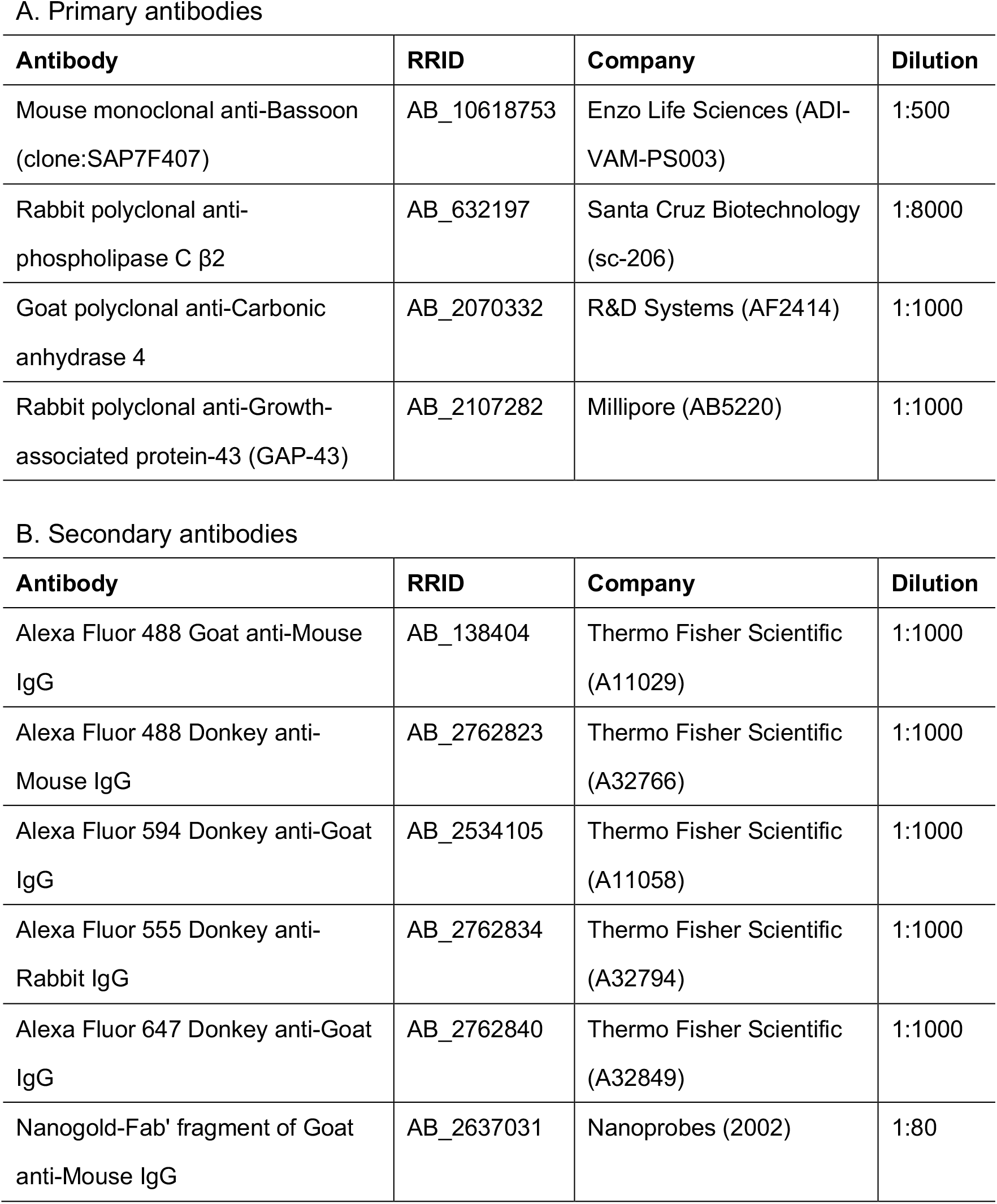

The antibody to Bassoon, SAP7F407, was raised against a glutathione S-transferase-fusion protein containing amino acid residues 756–1001 of rat Bassoon. This antibody recognizes a single band of ∼420 kilodaltons in an immunoblot analysis of a rat brain tissue lysate (manufacturer’s data sheet). The immunoreactivity (IR) of SAP7F407 was previously reported at presynaptic sites in the retina (Dick, et al., 2003), cerebellum and hippocampus (Richter et al., 1999), and the calyx of Held (Dondzillo, et al., 2010). SAP7F407-IR in the retina was shown to be eliminated in *Bassoon* gene-targeted mice (Dick, et al., 2003).

### Immunoelectron microscopy

The detection of the Bassoon-IR by a pre-embedding immunogold method was performed as previously described (Ikuta, et al., 2021). Tissues were cut into 10-µm-thick sections on a cryostat, collected on silane-coated slides, and air-dried for 15 min. After rinsing with PBS, sections were blocked with PBS containing 20% Blocking One and 0.005% saponin for 15 min. Sections were incubated with the anti-Bassoon antibody at 4°C for 42 - 48 hr. Negative controls were prepared by omitting the primary antibody. After rinsing with PBS containing 0.005% saponin, sections were incubated with the Fab’ fragment of goat anti-mouse IgG labeled with 1.4 nm colloidal gold (Nanoprobes, NY, USA)(Table 1) at 4°C overnight. After rinsing four times with PBS containing 0.005% saponin for 10 min each, sections were further rinsed with 0.1 M PB. Sections were then fixed with 1% glutaraldehyde in 0.1 M PB for 10 min. After rinsing with 0.1 M PB, sections were rinsed with 50 mM HEPES (pH 5.8) three times for 5 min each. The binding sites of the gold-labeled antibody were silver-enhanced using a kit (HQ Silver, Nanoprobes), followed by two washes with water. Sections were post-fixed with 0.5% osmium tetroxide in 0.1 M PB at 4°C for 1.5 hr. After rinsing with water, sections were dehydrated by passing through a graded series of ethanol and propylene oxide, and then embedded in epoxy resin. Ultrathin sections were cut on an ultramicrotome, collected on grids, stained with uranyl acetate and lead citrate, and observed under an electron microscope (JEM-1400 plus, JEOL, Japan). The contrast of images was adjusted with Adobe Photoshop.

## Results

Using confocal laser microscopy with immunofluorescence, we examined the localization of Bassoon in the taste buds of the circumvallate papillae of mice. Bassoon-IR was detected as round or elongated puncta (Figure 1). In longitudinal sections of the taste buds, a large number of Bassoon-positive puncta were observed in a line (Figure 1d).

**Figure 1.**
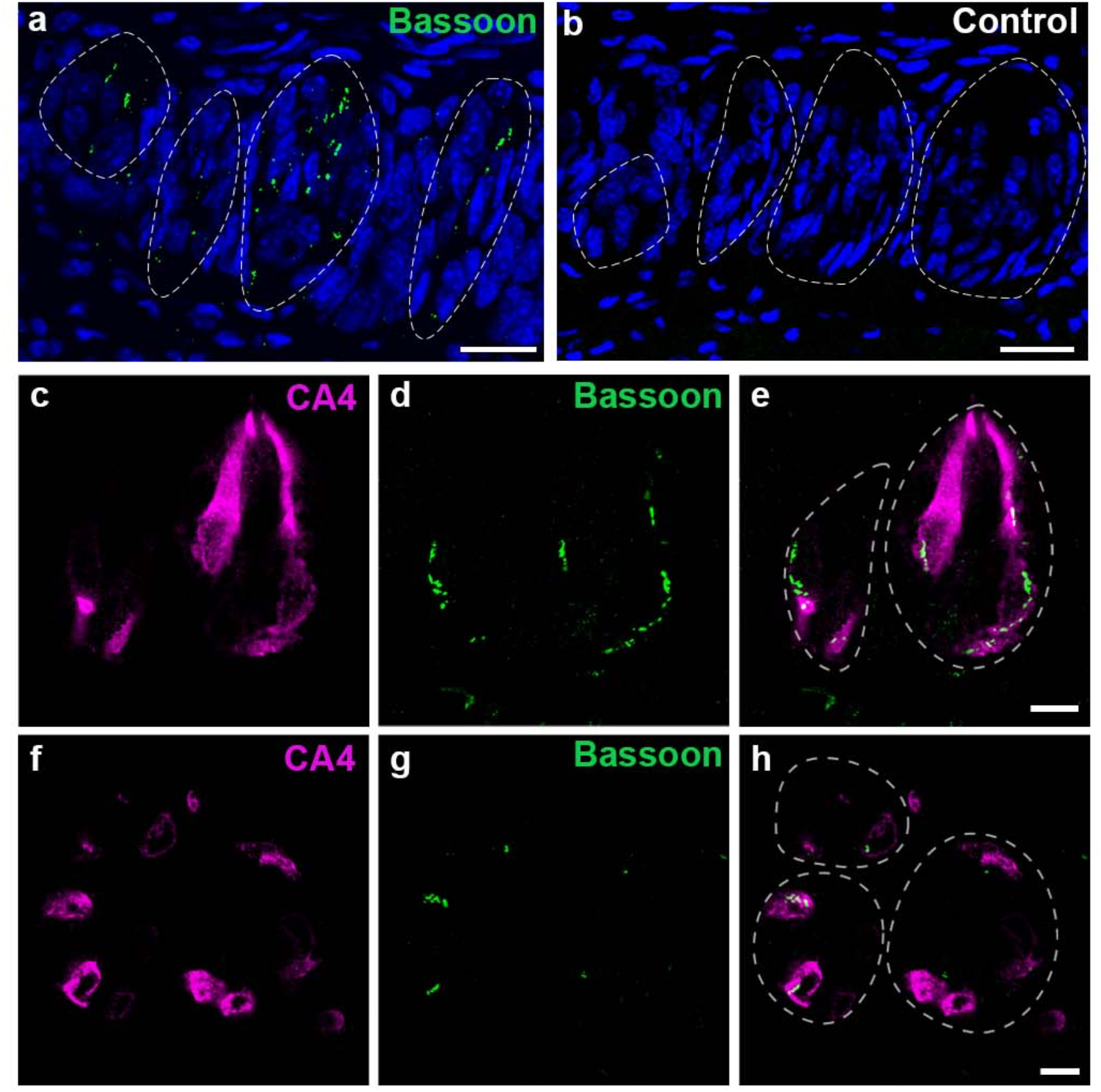
Representative confocal images (individual Z sections) of single (a) or double immunolabeling for Bassoon and the Type III cell marker, CA4 (c-h). Bassoon was detected as round or elongated puncta in taste buds (a, d and g). No immunolabeling was observed in the negative control (b). Nuclei were stained with DAPI (a, b). Double immunolabeling for Bassoon and CA4 showed that most Bassoon-positive puncta were observed in CA4-positive cellular profiles (c-h). Longitudinal (c-e) and transverse (f-h) images to the long axis of taste buds. Scale bars = 20 μm (a, b); 10 μm (e, h).

Since Bassoon is a presynaptic protein of conventional synapses, which are observed between Type III cells and nerve fibers in taste buds, its localization to Type III cells was expected. To confirm this, we performed double immunolabeling for Bassoon and CA4, a marker for Type lll cells (Chandrashekar, et al., 2009). As expected, most Bassoon-IR was observed in CA4-positive cellular profiles (Figure 1c-h). We investigated whether Bassoon localized to Type ll cells, the other taste receptor cells. We used PLCβ2 as a Type ll cell marker (Clapp, et al., 2004), and found that few PLCβ2-positive profiles contained Bassoon-IR (5 of 478 Type II cells, n = 3), while approximately 50% of the CA4-positive profiles observed contained Bassoon-IR (84 of 169 Type III cells, n = 3) (Figure 2a-e).

**Figure 2.**
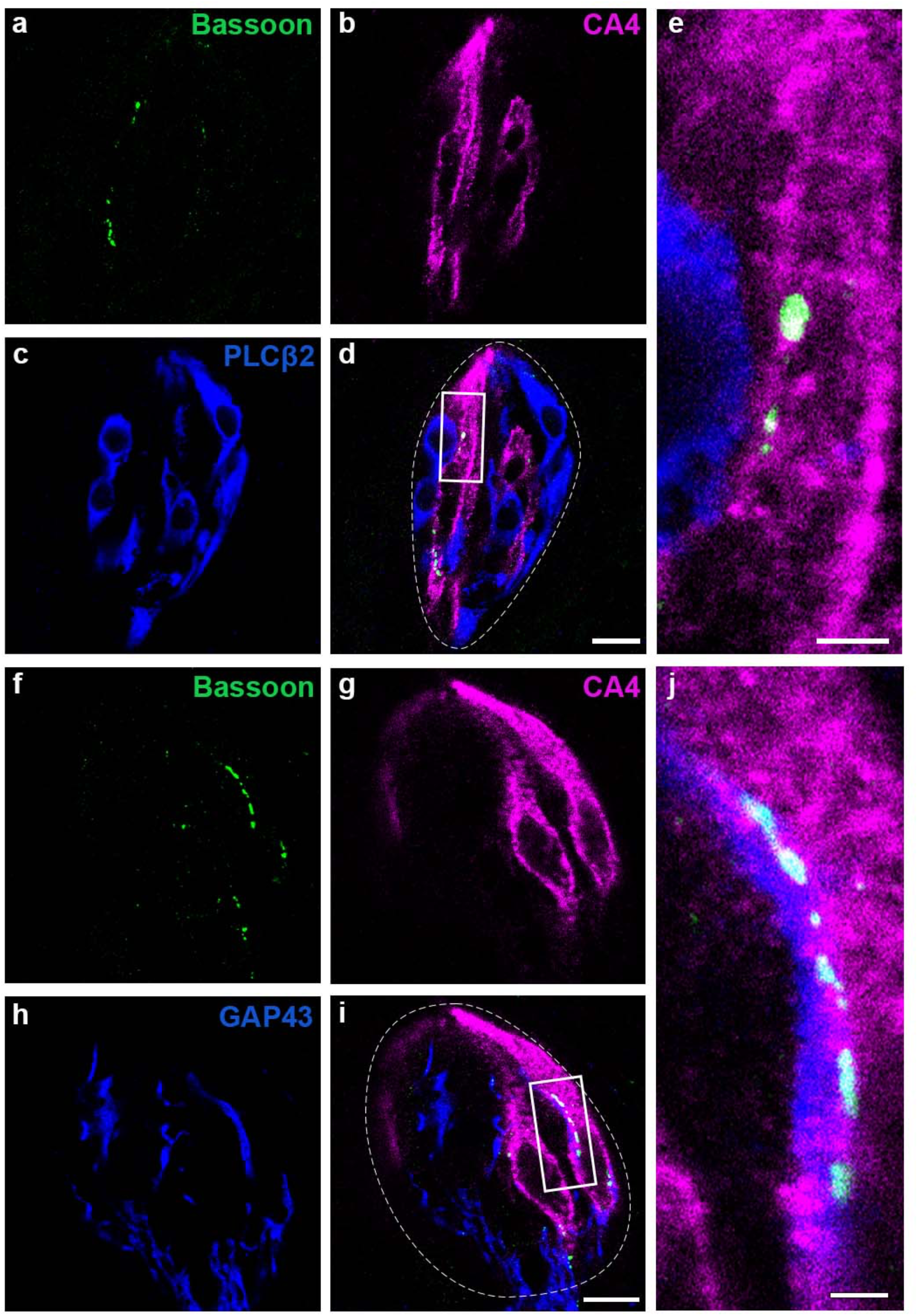
Representative confocal images (individual Z sections) of triple immunolabeling for Bassoon, CA4 and the Type II cell marker, PLCβ2 (a-e), or Bassoon, CA4 and the nerve fiber marker, GAP-43 (f-j). Triple labeling for Bassoon (a), CA4 (b), and PLCβ2 (c) showed that Bassoon-positive puncta were detected in CA4-positive cells but rarely in PLCβ2-positive cells (d, e). Panel e corresponds to the rectangle area in d. Triple labeling for Bassoon (f), CA4 (g) and GAP-43 (h) showed that Bassoon-positive puncta were observed in the regions of CA4-positive cells contact with nerve fibers (i, j). Panel j corresponds to the rectangle area in i. Scale bars = 10 μm (d, i); 2 μm (e, j).

We then examined the spatial relationship between Bassoon-IR and intragemmal nerve fibers. We used GAP-43 as a marker for intragemmal nerve fibers because the distribution pattern of GAP-43-IR in taste buds closely overlapped with that of P2X2 receptor-IR (Zaidi, et al., 2016), which is also used as a marker for intragemmal nerve fibers. Triple immunolabeling for GAP-43, CA4, and Bassoon showed that the majority of Bassoon-IR was in proximity to or partially overlapping with both GAP43-positive fibers and CA4-positive cellular profiles (Figure 2f-j).

We used a pre-embedding immunogold method with silver enhancement to examine the localization of Bassoon in taste buds at the ultrastructural level (Figure 3). Most of the cells showing Bassoon-IR had nuclear profiles with moderate chromatin densities among taste bud cells, and frequently showed nuclear envelope invaginations (Figure 3a). These characteristics of nuclei are consistent with those of Type III cells (Takeda and Hoshino, 1975; Yang et al., 2020). Bassoon-IR was not detected in Type II cells, which are electron-translucent cells that have oval or round nuclei with an abundant cytoplasm (Murray, 1993;Yang et al., 2020).

**Figure 3.**
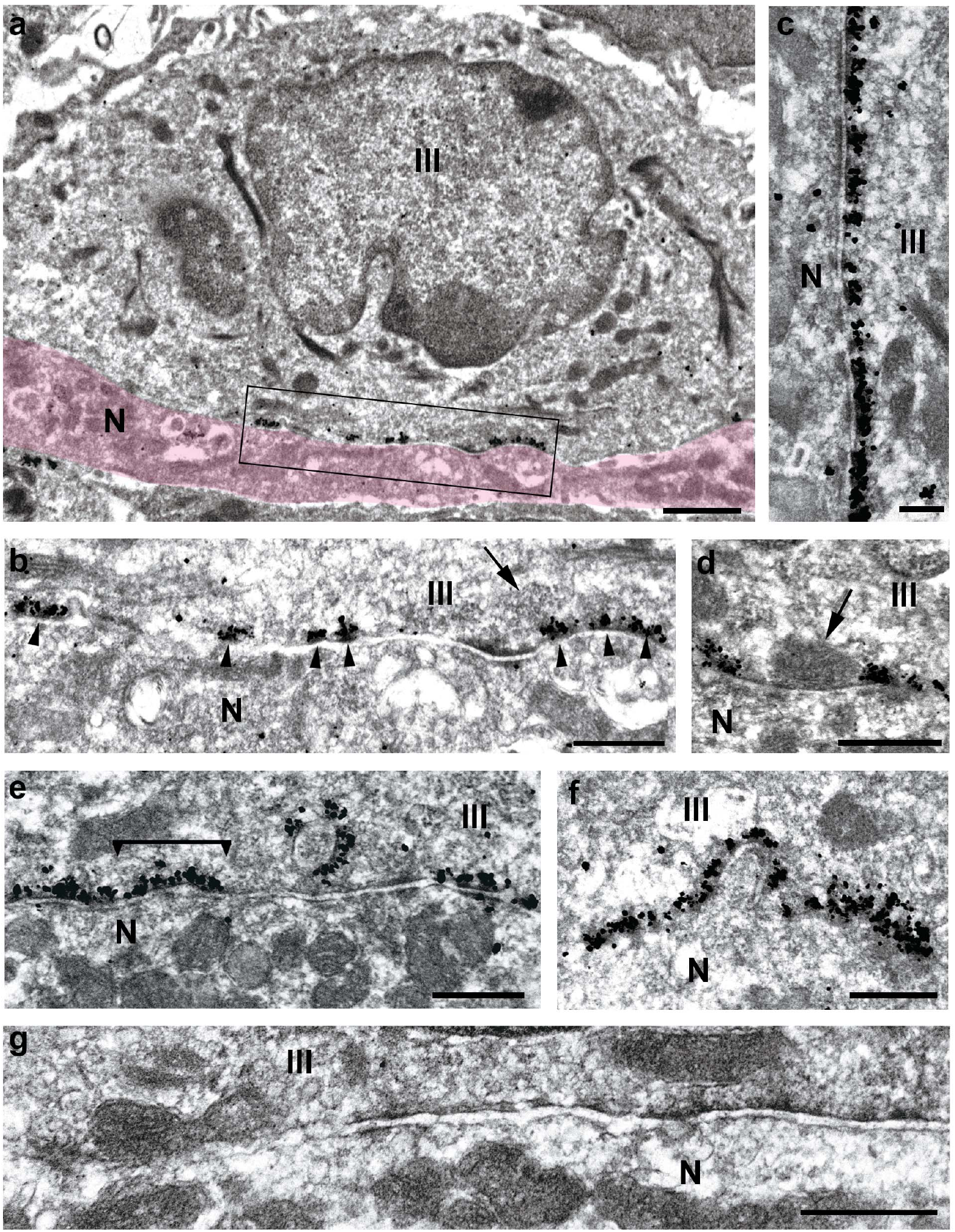
Representative immunoelectron micrographs showing the localization of Bassoon-IR in taste buds. Silver deposits for Bassoon-IR were observed along the plasma membranes of Type lll cells apposed to nerve fibers. The region labeled in red is a profile of an afferent nerve fiber (a). A higher magnification of the boxed area in a is shown in b. Arrowheads indicate clusters of silver deposits. Some of the clusters of Bassoon-IR were accompanied by clusters of synaptic vesicles (b, arrow; e, bar). The membrane thickening of nerve fibers was frequently observed at the regions apposed to the membranes of Type III cells bearing Bassoon-IR (c, e). Bassoon-IR was also observed around mitochondria (d, arrow) that adhered to the plasma membranes apposed to nerve fibers (d). Bassoon-IR was observed at the active zones of “fingerlike” synapses (e, f). Few silver deposits were observed in the negative control (g). Type IIl cells and afferent nerve fibers are labeled with “lll” and “N,” respectively. Scale bars = 1 μm (a); 500 nm (b, d–g); 250 nm (c).

The silver deposits of Bassoon-IR were observed along the inner side of the membranes of Type III cells apposed to the nerve fibers. Silver deposits appeared to cluster in an intermittent manner at the membrane regions at which Bassoon-IR localized (Figure 3b, c). Some of the clusters of silver deposits were accompanied by clusters of synaptic vesicles (Figure 3b, e). These silver deposit clusters were often observed in groups along the plasma membrane of Type III cells in contact with nerve fibers. The groups of clusters of Bassoon-IR sometimes extended to several micrometers (Figure 3c). This distribution pattern of Bassoon-IR is consistent with that of the active zones of conventional synapses in taste buds (Royer and Kinnamon, 1988). In contrast to the abundant silver deposits in Type III cells, few were observed in the afferent nerve fibers in contact with Type III cells. The membrane thickening of nerve fibers was frequently noted at the regions apposed to the membranes of Type III cells bearing clusters of silver deposits (Figure 3c, e). Furthermore, the intercellular spaces between the membranes of Type III cells bearing Bassoon-IR and those of nerve fibers were filled with an amorphous substance (Figure 3c, e). These results indicated that silver deposits localized at the active zones of synapses in Type III cells.

We also detected Bassoon-IR around mitochondria that tightly adhered to the plasma membranes apposed to nerve fibers (Figure 3d). This Bassoon-IR around mitochondria was considered to correspond to the active zones of synaptic ensembles (Royer and Kinnamon, 1988) or mixed synapses (Yang et al., 2020).

We occasionally observed circular membrane structures with Bassoon-IR in Type III cells (Figure 3e). These structures with Bassoon-IR were likely to be “fingerlike” synapses (Kinnamon et al., 1985;Royer and Kinnamon, 1988), which are characterized by a protrusion of the nerve fiber into an invagination of Type III cells. The profiles of the active zones of fingerlike synapses vary in shape from invaginations to circles depending on the planes of ultrathin sections. Figure 3f shows the distribution pattern of Bassoon-IR in a fingerlike synapse shown as a simple invagination.

## Discussion

Bassoon-IR was mostly observed in mouse taste buds as puncta in CA4-positive cellular profiles, and was detected in the vicinity of intragemmal nerve fibers at the light microscopic level. At the ultrastructural level, the distribution pattern of Bassoon-IR was in accordance with that of the active zones of conventional synapses in the taste buds, including fingerlike synapses and synapses with mitochondria that adhered to the presynaptic membranes. These results indicate that Bassoon in the mouse taste buds almost exclusively localized to the active zones of conventional synapses between Type III cells and afferent nerve fibers.

The localization of several synaptic proteins, including synapsin I (Finger, et al., 1990), SV2 (Pumplin and Getschman, 2000), synaptophysin (Asano-Miyoshi, et al., 2009;Pumplin and Getschman, 2000), synaptogyrin-1 (Kotani, et al., 2013), synaptobrevin/VAMP2 (Pumplin and Getschman, 2000;Yang, et al., 2004), synaptotagmin-1 (Kohno, et al., 2005), syntaxin-1 (Pumplin and Getschman, 2000;Yang, et al., 2007), and SNAP-25 (Pumplin and Getschman, 2000;Yang, et al., 2000), has been examined in taste buds; however, none exclusively localized to synaptic sites. These synaptic proteins were all detected in intragemmal nerve fibers. Furthermore, synaptogyrin-1, synaptobrevin/VAMP2, and syntaxin-1 were found in both Type II and Type III cells. Although synaptotagmin-1 and SNAP-25 were only observed in Type III cells, they were diffusely detected in the cytoplasm and were not concentrated at synaptic sites. Therefore, none of these synaptic proteins are markers for conventional synapses between Type III cells and nerve fibers. In contrast to the synaptic proteins previously examined in taste buds, the present results indicate that Bassoon is a reliable marker for synapses between Type III cells and afferent nerve fibers in taste buds. Immunohistochemistry for Bassoon will contribute to investigations on the formation of and impairments in conventional synapses in taste buds.

## Ethical approval

The care and use of animals were approved by the Institutional Animal Care and Use Committee of Fukuoka Women’s University in compliance with Guidelines for Proper Conduct of Animal Experiments (Science Council of Japan).

